# Intramolecular quality control: HIV-1 Envelope gp160 signal-peptide cleavage as a functional folding checkpoint

**DOI:** 10.1101/2020.07.08.188672

**Authors:** Nicholas McCaul, Matthias Quandte, Ilja Bontjer, Guus van Zadelhoff, Aafke Land, Rogier W. Sanders, Ineke Braakman

## Abstract

Removal of the membrane-tethering signal peptides that target secretory proteins to the endoplasmic reticulum is a prerequisite for proper folding. While generally thought to be removed well before translation termination, we here report two novel post-targeting functions for the HIV-1 gp120 signal peptide, which remains attached until gp120 folding triggers its removal. First, the signal peptide improves fidelity of folding by enhancing conformational plasticity of gp120 by driving disulfide isomerization through a redox-active cysteine, at the same time delaying folding by tethering the N-terminus to the membrane, which needs assembly with the C-terminus. Second, its carefully timed cleavage represents intramolecular quality control and ensures release and stabilization of (only) natively folded gp120. Postponed cleavage and the redox-active cysteine both are highly conserved and important for viral fitness. Considering the ∼15% secretory proteins in our genome and the frequency of N-to-C contacts in protein structures, these regulatory roles of the signal peptide are bound to be more common in secretory-protein biosynthesis.

## Introduction

The endoplasmic reticulum (ER) is home to a wealth of resident chaperones and folding enzymes that cater to approximately a third of all mammalian proteins during their biosynthesis (Ellgaard et al., 2016; Kanapin et al., 2003). It is the site of N-linked glycan addition and disulfide-bond formation, both of which contribute to protein folding, solubility, stability, and function. Targeting to the mammalian ER in general is mediated by N-terminal signal peptides, which direct the ribosome-nascent chain complex to the membrane and initiate co-translational translocation (Blobel and Dobberstein, 1975; Gorlich et al., 1992; Görlich et al., 1992; Jackson and Blobel, 1977; Lingappa et al., 1977; Walter, 1981). For soluble and type-I transmembrane proteins, the N-terminal signal peptide is 15-50 amino acids long and contains a cleavage site recognized by the signal peptidase complex (von Heijne, 1985). While a great deal of sequence variation occurs between signal sequences, conserved features do exist. These include a positively charged, N-terminal n-region, a hydrophobic h-region and an ER-lumenal c-region (von Heijne, 1983, 1984, 1985).

Classic paradigm-establishing studies showed that cleavable signal peptides are removed co-translationally, immediately upon exposure of the cleavage site in the ER lumen (Blobel and Dobberstein, 1975; Jackson and Blobel, 1977). This would imply that signal peptides function only as cellular postal codes and that signal-peptide cleavage and folding are independent events. Evidence is emerging however that increased nascent-chain lengths are required for cleavage (Daniels et al., 2003; Hegde and Bernstein, 2006; Rutkowski et al., 2003), indicating that the signal peptidase does not cleave each consensus site immediately upon translocation into the ER lumen. Examples are the influenza-virus hemagglutinin, in which signal-peptide cleavage occurs on the longer nascent chain, well after glycosylation (Daniels et al., 2003), EDEM1 (Tamura et al., 2011), human cytomegalovirus (HCMV) protein US11 (Rehm et al., 2001), and HIV-1 envelope glycoprotein gp160 (Li et al., 1994, 2000), suggesting that signal peptides can function as more than mere postal codes. Late signal-peptide cleavage is easily overlooked because Western blots lack temporal resolution and may mask small mass differences.

Gp160 is the sole antigenic protein on the surface of the HIV-1 virion and mediates HIV-1 entry into target cells (Wyatt and Sodroski, 1998). It folds and trimerizes in the ER, leaves upon release by chaperones and packaging into COPII-coated vesicles, and is cleaved by Golgi furin proteases into two non-covalently associated subunits: the soluble subunit gp120 (Figure 1A, in colors), which binds host-cell receptors, and the transmembrane subunit gp41 (Figure 1A, uncolored), which contains the fusion peptide (Decroly et al., 1994; Earl et al., 1990; Earl et al., 1991; Hallenberger et al., 1992; Wyatt and Sodroski, 1998). The so-called outer-domain residues [according to (Pancera et al., 2010)] are colored in pink (Figure 1A), the inner domain, which folds from more peripheral parts of the gp120 sequence, in grey, and the variable loops in green. Correct function of gp160 requires proper folding including oxidation of the correct cysteine pairs into disulfide bonds (Bontjer et al., 2009; Land and Braakman, 2001; Land et al., 2003; Sanders et al., 2008; Snapp et al., 2017). Disulfide-bond formation and isomerization in gp160 begin co-translationally, on the ribosome-attached nascent chain, and continue long after translation, until the correct set of ten conserved disulfide bonds have been formed (Land and Braakman, 2001; Land et al., 2003). The soluble subunit gp120 can be expressed independently of gp41 and folds with highly similar kinetics as gp160 (Land et al., 2003). Signal-peptide cleavage only occurs once gp120 attains a near-native conformation and requires both N-glycosylation and disulfide-bond formation, but is gp41 independent (Land et al., 2003; Li et al., 1996). Mutation-induced co-translational signal-peptide cleavage changes the folding pathway of gp120 and is disadvantageous for viral function (Pfeiffer et al., 2006; Snapp et al., 2017). Given the interplay between signal-peptide cleavage and gp120 folding, we set out to investigate the mechanism that drives post-translational cleavage and its relevance for gp120 folding and viral fitness.

**Figure 1.**
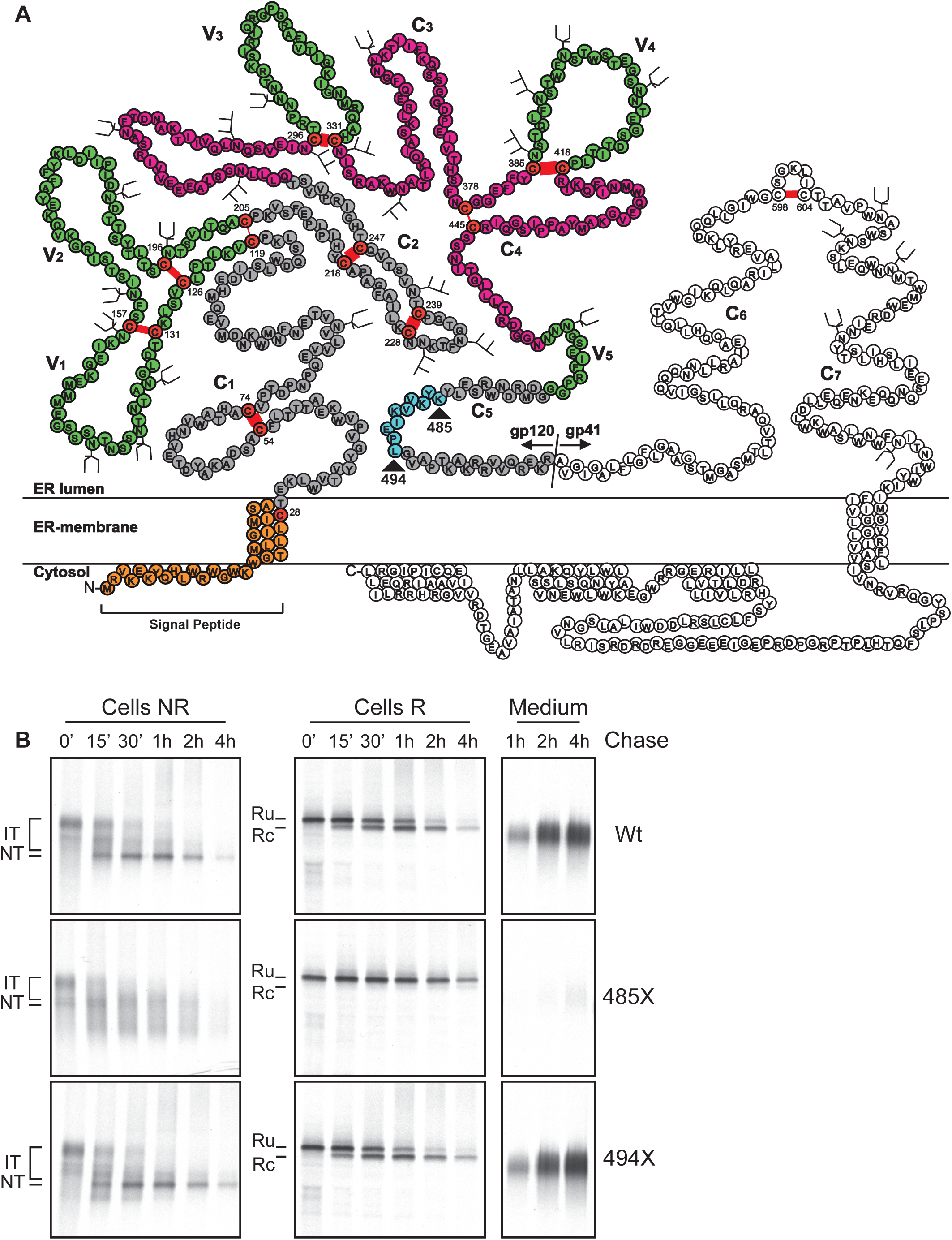
Signal-peptide cleavage requires the gp120 C-terminus. **A)** Schematic representation of gp160 amino-acid sequence with its signal peptide (orange) still attached [adapted from (Leonard et al., 1990)]. Gp120 inner domain (grey) and outer domain (pink) [according to (Pancera et al., 2010)], cysteines (red) are numbered and disulfide bonds represented by red bars. Thickness of disulfide bonds is representative of their importance for folding and (or) infectivity [thickest essential for folding, middle dispensable for folding, essential for infectivity, thinnest dispensable for both folding and infectivity (van Anken et al., 2008)]. Gp120 contains five constant regions (C1-C5) and five variable regions (green, V1-V5). Oligomannose and complex glycans are represented as three- or two-pronged forked symbols respectively [adapted from (Leonard et al., 1990)]. Amino-acid stretch 485-494 is marked in teal. **B)** HeLa cells transiently expressing gp120 Wt and C-terminal truncations were radiolabeled for 5 minutes and chased for the indicated times. After detergent lysis, samples were immunoprecipitated with polyclonal antibody 40336. After immunoprecipitation, samples were analyzed by non-reducing (Cells NR) and reducing (Cells R) 7.5% SDS-PAGE. Gp120 was immunoprecipitated from medium samples with antibody 40336 and directly analyzed by reducing 7.5% SDS-PAGE (Medium). Gels were dried and exposed to Kodak-MR films or Fujifilm phosphor screens for quantification. Ru: reduced, signal peptide cleaved gp120, Rc: reduced, signal-peptide-uncleaved gp120, IT: intermediates, NT: native.

We used various kinetic oxidative-folding assays on gp120 combined with functional studies on recombinant HIV strains encoding gp160 mutants, and discovered a novel role for the ER-targeting signal peptide as quality-control checkpoint and folding mediator. A conserved cysteine in the membrane-tethered signal peptide drives disulfide isomerization in the gp120 ectodomain until gp120 folding triggers signal-peptide cleavage and release of the N-terminus. We uncovered this functional, mutual regulation as an intramolecular quality control that ensures native folding of a multidomain glycoprotein.

## Results

### Signal-peptide cleavage requires the gp120 C-terminus

Of the nine disulfide bonds in gp120, five are critical for proper folding and signal-peptide cleavage, three in the constant regions of gp120 in the (grey) inner domain, and two in the outer (pink) domain at the base of variable loops (green) V3 and V4 [Figure 1A, (van Anken et al., 2008)]. Gp120 undergoes extensive disulfide isomerization during its folding process as seen from the smear of gp120 folding intermediates (IT) in the non-reducing gel upon ^35^S-radiolabeling, from reduced gp120 down to beyond native gp120 [(Land et al., 2003); Figure 1B, NR, 0’ chase]. This smear gradually disappears into a native band with discrete mobility, at around the time the signal peptide is removed (Figure 1B R, from 15’ chase). Yet, it is far from obvious which aspect of gp120 folding triggers signal-peptide cleavage. We therefore embarked on the linear approach and prepared C-terminal truncations of gp120 from 110-aa length (111X, a gp120 molecule truncated after position 110) to full-length and analyzed in which the signal peptide was cleaved (Figure S1). Radioactive pulse-chase experiments showed that only full-length gp120 (511 residues long) and 494X lost their signal peptides, but that in all shorter forms, including 485X, the signal peptide remained uncleaved and hence attached to the protein (Figure S1). We continued with a time course for 485X and 494X to examine and compare their folding pathways (Figure 1B). Both truncations encompass the entire gp120 sequence except for the last 26 and 17 amino acids, respectively (Figure 1A).

Like wild-type gp120, immediately after pulse labeling (synthesis) the 2 C-terminally truncated mutants ran close to the position of reduced gp120 in non-reducing SDS-PAGE. The 485X truncation formed disulfide bonds towards a compact structure, as the folding intermediates IT ran lower in the gel than reduced protein. It failed to form native gp120 however (NT, Figure 1B) or another stable intermediate. Instead it acquired compactness far beyond the mobility of NT and remained highly heterogeneous, suggesting the formation of long-distance disulfide bonds that increased compactness and hence electrophoretic mobility (Figure 1B, Cells NR, 15’-2h). Only a fraction, if any, of the 485X mutant lost its signal peptide or acquired competence to leave the ER and be secreted (Figure 1B, Medium), even though all cysteines were present in the 485X gp120 protein. In contrast, addition of only 9 residues made 494X behave like wild-type gp120: cleavage rate and secretion were indistinguishable (Figure 1B). Oxidative folding progressed similarly as well, except for a transient non-native disulfide-linked population that ran more compact than NT (Figure 1B, Cells NR, 0-1 h) and disappeared over time (Cells NR, 2-4 h). We concluded that signal-peptide cleavage required synthesis and folding of more than 484 out of the 511 amino acids of gp120. The downstream amino acids in the C5 region in the inner (grey) domain (Figure 1A, teal) triggered the switch from non-cleavable to cleavable.

### A pseudo salt bridge in the inner-domain β-sandwich controls signal-peptide cleavage and gp160 function

Amino acids 485-494 form a β-strand (Figures 2A and 2B, teal, β31), which is part of the β-sandwich in the inner (grey) domain of gp120 (Figures 1A, 2A, and 2B) [coding of strands and helices from (Garces et al., 2015)]. This β-sandwich is formed by interactions of seven β-strands in constant domains C1, C2, and C5 (Figures 1A, 2A, and 2B). Six of the strands are close to the N-terminus and the 6^th^ strand, β31 (teal), is contributed by the C-terminal C5 region. As the addition of β31 triggered signal-peptide removal we hypothesized that the complete and properly folded β-sandwich was the minimal requirement for cleavage.

**Figure 2.**
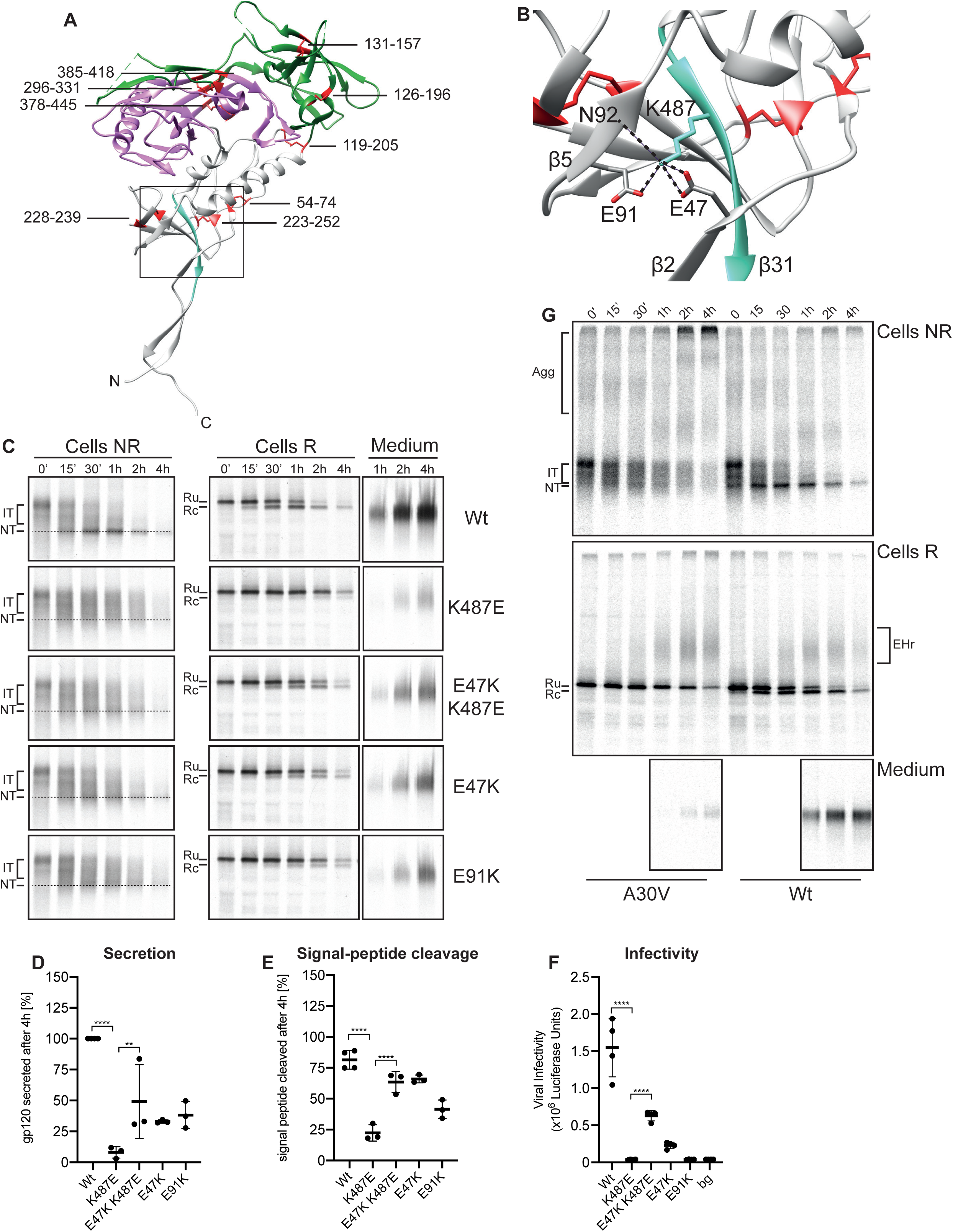
Integration of gp120 N- and C-terminus regulates signal-peptide cleavage. **A)** Gp120 crystal structure, 5CEZ (Garces et al., 2015), domains are colored as in Figure 1A. N and C termini are indicated; disulfide bonds are shown as red lines. Inner-domain β-sandwich is boxed. **B)** Zoom in of inner-domain β-sandwich. C-terminal β-strand in teal with K487 forming hydrogen bonds (dashed lines) with E47, E91 and main-chain oxygen of N92. Beta strands are numbered, and disulfide bonds indicated as red lines. Amino acids are named and numbered according to HXB2 sequence. **C)** Experiments as in Figure 1C with HeLa cells expressing Wt gp120 or indicated β-sandwich mutants. **D)** Quantifications of experiments performed as in C, intracellular levels at 15’ were used to correct for differences in expression between mutants and corrected values compared to wild-type secretion at 4 h. Error bars: SD. **E)** As in **D** except % signal peptide cleaved at 4 h was measured from reducing gels. **F)** Luciferase-based infectivity assay on TZM-bl cells. Cells were infected with 100 pg of HIV-1 LAI virus containing Wt or mutant gp160. Error bars: SD. **G)** Pulse-chase performed as in Figure 1B. **: p<0.01, ****: p<0.0001. Complete statistical values are listed in Table 1.

**Table 1.**
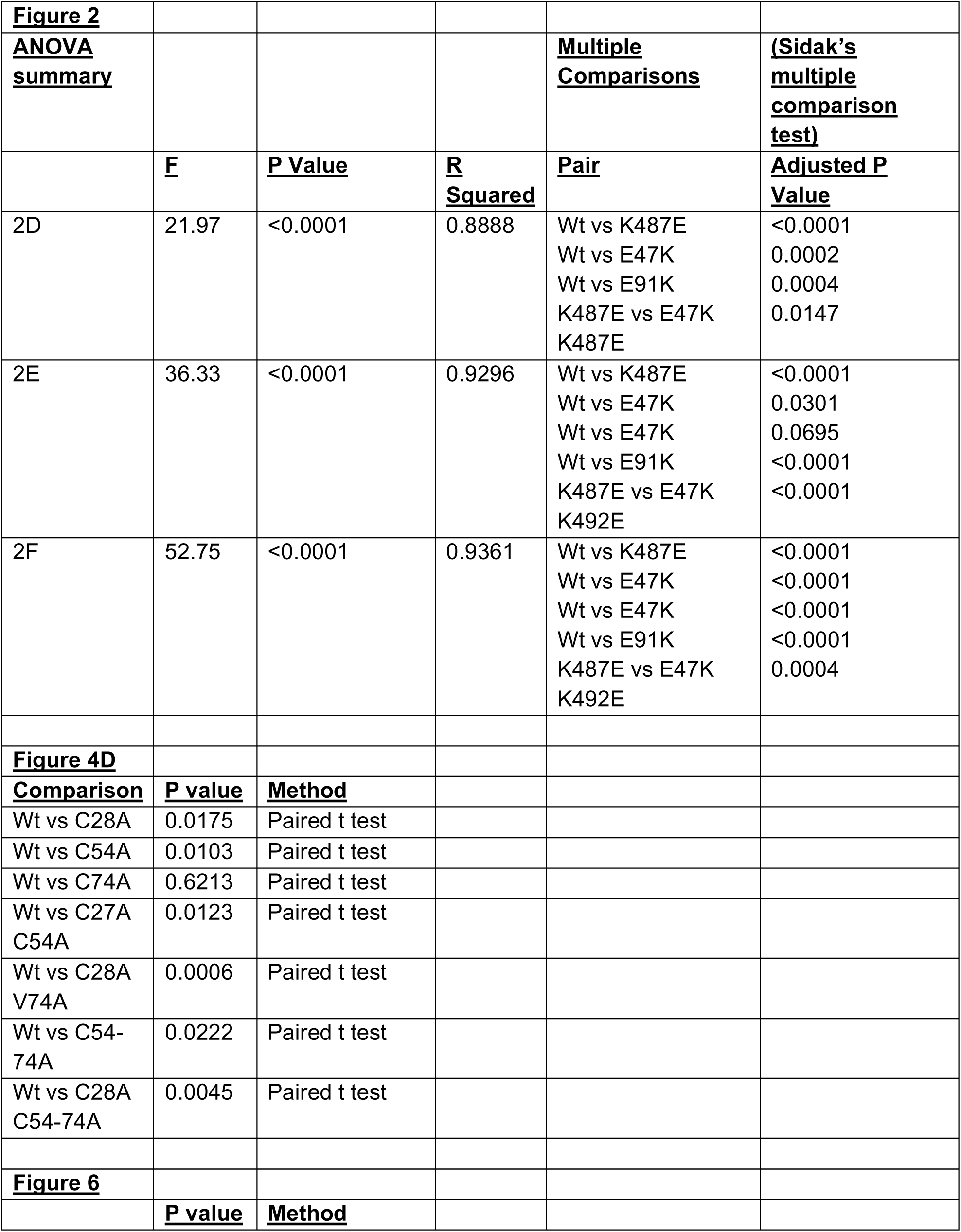
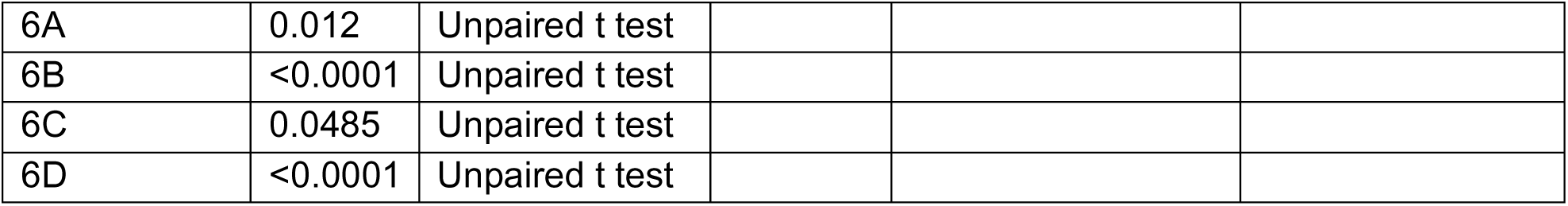
Complete statistical reporting for experiments in Figures 2, 4 and 6.

|  |  |  |  |  |  |
| --- | --- | --- | --- | --- | --- |
| **<u>Figure 2</u>** |  |  |  |  |  |
| **<u>ANOVA summary</u>** |  |  |  | **<u>Multiple Comparisons</u>** | **<u>(Sidak's multiple comparison test)</u>** |
|  | **<u>F</u>** | **<u>P Value</u>** | **<u>R Squared</u>** | **<u>Pair</u>** | **<u>Adjusted P Value</u>** |
| 2D | 21.97 | <0.0001 | 0.8888 | Wt vs K487E<br>Wt vs E47K<br>Wt vs E91K<br>K487E vs E47K<br>K487E | <0.0001<br>0.0002<br>0.0004<br>0.0147 |
| 2E | 36.33 | <0.0001 | 0.9296 | Wt vs K487E<br>Wt vs E47K<br>Wt vs E47K<br>Wt vs E91K<br>K487E vs E47K<br>K492E | <0.0001<br>0.0301<br>0.0695<br><0.0001<br><0.0001 |
| 2F | 52.75 | <0.0001 | 0.9361 | Wt vs K487E<br>Wt vs E47K<br>Wt vs E47K<br>Wt vs E91K<br>K487E vs E47K<br>K492E | <0.0001<br><0.0001<br><0.0001<br><0.0001<br>0.0004 |
| **<u>Figure 4D</u>** |  |  |  |  |  |
| **<u>Comparison</u>** | **<u>P value</u>** | **<u>Method</u>** |  |  |  |
| Wt vs C28A | 0.0175 | Paired t test |  |  |  |
| Wt vs C54A | 0.0103 | Paired t test |  |  |  |
| Wt vs C74A | 0.6213 | Paired t test |  |  |  |
| Wt vs C27A<br>C54A | 0.0123 | Paired t test |  |  |  |
| Wt vs C28A<br>V74A | 0.0006 | Paired t test |  |  |  |
| Wt vs C54-74A | 0.0222 | Paired t test |  |  |  |
| Wt vs C28A<br>C54-74A | 0.0045 | Paired t test |  |  |  |
| **<u>Figure 6</u>** |  |  |  |  |  |
|  | **<u>P value</u>** | **<u>Method</u>** |  |  |  |

|  |  |  |
| --- | --- | --- |
| 6A | 0.012 | Unpaired t test |
| 6B | <0.0001 | Unpaired t test |
| 6C | 0.0485 | Unpaired t test |
| 6D | <0.0001 | Unpaired t test |

To address this, we designed charge mutants aimed to prevent assembly of β31 with the N-terminal part of the β-sandwich (Figure 2B). In the high-resolution crystal structure (Garces et al., 2015) K487 (in β31) forms hydrogen bonds with E47 (in β2), E91 (in β5), and the main-chain oxygen of N92 (in β5) (Figure 2B). We created charge-reversed mutants of the N-terminal glutamates (E47K and E91K), the C-terminal lysine (K487E) and combinations thereof (Figures 2C and S2). We did not include N92 in our mutagenesis study since its interaction involves the main-chain oxygen, which cannot be removed; we considered this inappropriate for our question.

As gp120 is the dominant subunit in gp160 folding and signal-peptide cleavage, and allows more detailed analysis because it is smaller than gp160, we subjected wild-type gp120 and all mutants to pulse-chase analysis of their oxidative folding, signal-peptide cleavage, and secretion (Figures 2C-E and S2A-D). Like the C-terminal truncations (485X, Figure 1B), all charge mutants in the β-sandwich formed gp120 molecules with higher electrophoretic mobility than native gp120 (NT), implying appreciable non-native long-range disulfide bonding, persisting at all time points or disappearing into aggregates (Figure S2B). K487E showed the strongest phenotype: minimal formation of NT and a much-delayed signal-peptide cleavage (Figures 2C, E and S2B, Cells NR and R, band Rc). This folding step was crucial for function as K487E mutant virus was non-infectious (Figure 2F). A striking rescue of the strong folding defect of K487E was effected by the charge reversal at the N-terminus: the double mutant E47K K487E displayed improved gp120 oxidation (Cells NR), signal-peptide cleavage (Cells R), and secretion (Medium) (Figures 2C-E and S2C), and rescued infectivity to ∼30% of wild type (Figure 2F). All N-terminal E-to-K mutants (E47K, E91K) oxidized and accumulated in a native-like position in gel but failed to form much of the sharp NT band seen in wild type (Figure 2C and S2A). The start of signal-peptide cleavage of both single mutants was delayed to 30-60 min after synthesis and total cleavage was 25-50% lower than wild type by 4 h (Cells R and Figure 2E and S2C). As a result, the secretion of all N-terminal mutants was decreased by ≥60% in 4 h compared to wild type (Medium, Figures 2C, D and S2A, C). The redundancy of two negative charges likely contributes to the intermediate folding phenotype of the N-terminal mutants and the ∼10% residual viral infectivity of the E47K mutant (Figure 2F).

We concluded that the C-terminal β-strand was essential for proper folding of the β-sandwich in the inner domain, which completes folding of gp120 and triggers signal-peptide cleavage. Timing of cleavage hence represents a checkpoint for proper folding of gp120.

### Retention of the signal peptide causes hypercompacting of gp160

During folding, gp120 undergoes extensive disulfide formation and isomerization before reaching its native state. These intermediates appear as “waves” on SDS-PAGE representing varying degrees of compactness of folding intermediates (Land et al., 2003). Because mutants of gp120 that exhibited delayed or absent cleavage all formed hypercompact forms that ran below NT (Figures 1B and 2C), we asked whether this heterogeneous electrophoretic mobility represented continued isomerization of gp120. We substituted the alanine in the -1-position relative to the cleavage site for a valine, to prevent cleavage by the signal-peptidase complex (Figure 2G, Cells R). Initial oxidative folding of A30V was similar to that of wild type (Figure 2G, NR). The A30V mutant, however, did not form a native band but populated all forms, from reduced to hypercompact oxidized, probably isomerizing continuously. The monomeric forms gradually disappeared into disulfide-linked, SDS-insoluble aggregates that increased in size and eventually became too large to enter the gel (Figure 2G, Agg).

In both wild-type and A30V gp120, an endoglycosidase H-resistant band appeared over time (Figure 2G, EHr). For wild-type gp120 this represents molecules that have transited through the Golgi complex and acquired an N-acetylglucosamine residue on their N-glycans but have yet to be secreted. For A30V gp120 this population may be due to the inaccessibility of some sugars for removal due to formation of SDS-insoluble aggregates.

We concluded that retention of the signal peptide either promotes formation of these hypercompact forms or prevents recovery from them. Because all signal-peptide-retaining mutants showed a high propensity of aggregation, it is likely that these SDS-insoluble aggregates are comprised of hypercompact forms of gp120. Tethering the N-terminus appears beneficial for folding, but release of gp120 from both tether and isomerization-driving cysteine is vital for stabilization of the acquired native fold and release from the ER.

### The cysteine in the signal peptide interacts with cysteines in gp160

As the signal peptide stays attached to gp160 for at least 15 min after chain termination it influences both co- and post-translational folding, through the tethering of the N-terminus to the membrane, as well as through interactions with the mature gp120 sequence (Snapp et al., 2017). The hypercompacting in non-cleaved mutants by continued disulfide isomerization (Figure 2G) implies that an unpaired cysteine must be available to keep attacking formed disulfide bonds. Opening once-formed disulfides may improve folding yield as the folding protein regains conformational freedom, and at the same time has a chance to recover from non-native disulfide bonding. Existing disulfides may be attacked by a cysteine from an oxidoreductase in the ER, or by an intramolecular cysteine in gp120 [as shown for BPTI (Weissman and Kim, 1992, 1993)]. The unpaired cysteine in position 28 within the signal peptide is a likely interaction candidate, because it is part of the consensus sequence for the signal peptidase and as such (partially) exposed to the ER lumen. During translocation, C28 may interact with gp160 cysteines while they pass through the translocon. Folding analysis as in Figures 1 and 2 however showed that mutating C28 had no detectable effect on oxidative folding (Figure 3A, C28A): folding intermediates disappeared, folded NT appeared, and the signal peptide was cleaved similarly and at similar times as wild-type gp120. Either C28A was identical to wild type or differences are missed due to asynchrony of the folding gp120 population. To amplify mobility differences, we alkylated with iodoacetic acid, which adds a charge to each free cysteine it binds to. To better synchronize the folding cohort, we modified the pulse-chase protocol with a preincubation with puromycin to release unlabeled nascent chains before labeling and added cycloheximide in the chase media to block elongation of radiolabeled nascent chains. At each chase time, gp120 C28A ran higher on a reducing gel (Figure S3), indicating that it has more free cysteines as it bound more iodoacetic acid than wild-type gp120. This could be due to either slower disulfide-bond formation, faster disulfide-bond reduction, or a combination of both, which suggests a role for C28 in the net gp120 disulfide formation or isomerization during folding.

**Figure 3.**
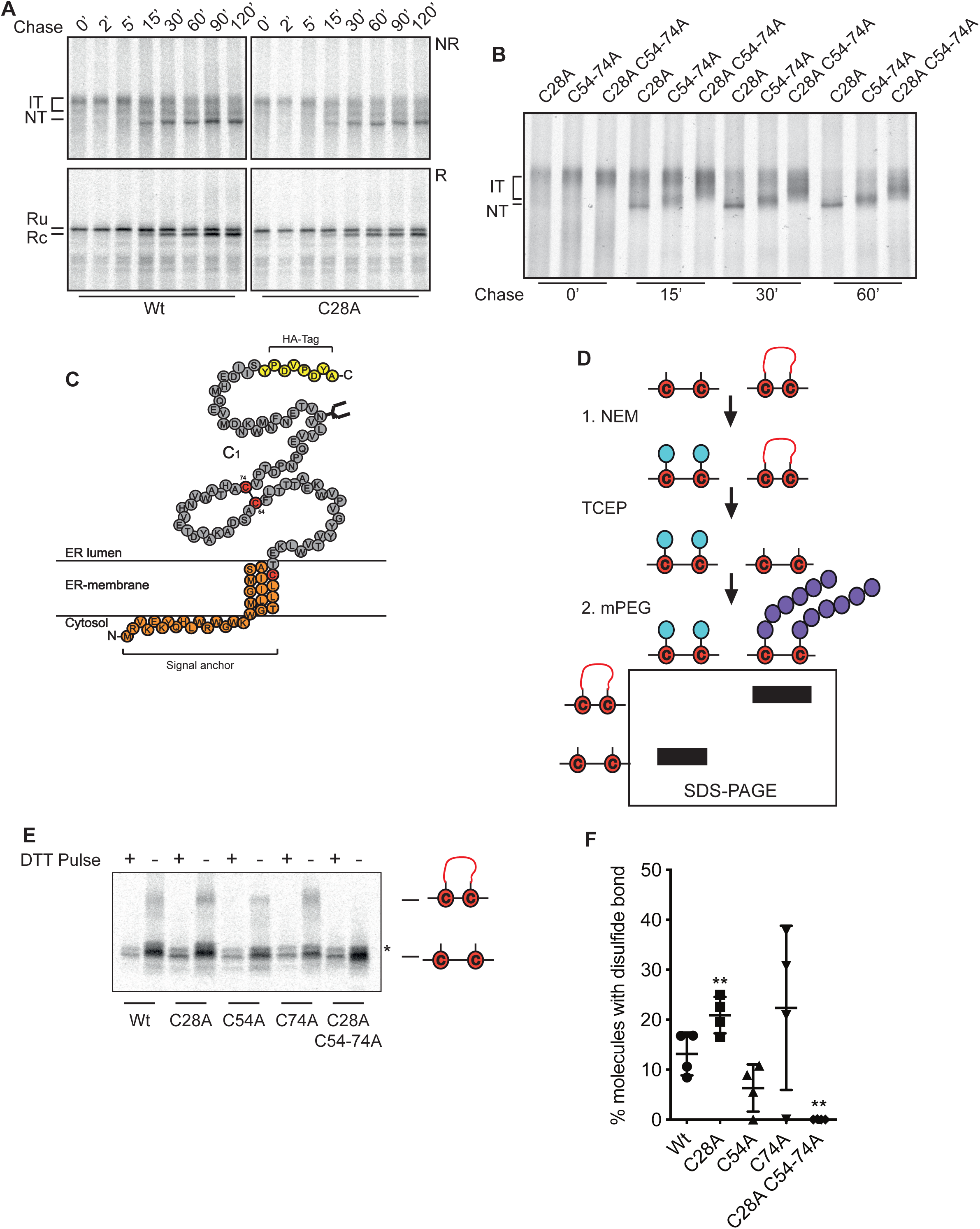
Signal-peptide cysteine is involved in gp120 oxidative folding. **A and B)** Pulse-chase experiments performed as in Figure 1B. Ru: reduced, signal peptide cleaved gp120, Rc: reduced, signal-peptide-uncleaved gp120, IT: intermediates, NT: native. **C)** Schematic representation of gp160 111X truncation construct with its signal anchor (orange), ectodomain (grey), numbered cysteines (red) and disulfide bond indicated by red bar; C-terminal HA tag in yellow; N-glycan depicted as forked structure. **D)** Schematic representation of mPEG alkylation-switch assay. In short, free cysteines are alkylated by NEM, which is excluded from disulfide bonds. Disulfide bonds are then reduced and resulting free cysteines are alkylated with mPEG-malemide which provides a 5 kDa shift per cysteine alkylated when analyzed by SDS-PAGE. **E)** HEK 293T cells expressing the indicated 111X truncations were pulse labeled for 30 minutes in the presence (+) or absence (–) of 5 mM DTT. At the end of the pulse, cells were scraped from dishes, homogenized and subjected to the double-alkylation mPEG-malemide alkylation protocol depicted in D (Appenzeller-Herzog and Ellgaard, 2008). After alkylation, samples were immunoprecipitated with a polyclonal antibody recognizing the HA-tag and analyzed by non-reducing 4-15% gradient SDS-PAGE. *: background band. **F)** Autoradiographs from experiments performed as described in E were quantified. Error bars: SD. **: p< 0.01; complete statistical values are listed in Table 1.

The importance of C28 became clear when we removed disulfide bond 54-74 in C1. Deletion of disulfide 54-74 prevents signal-peptide cleavage, but allows gp160 to reach a compact position just above natively folded protein NT [Figure 3B, (van Anken et al., 2008)]. When C28A was introduced into the 54-74 deletion, folding intermediates were blocked at a much earlier phase and remained significantly less compact (Figure 3B). The phenotype was the same when we combined C28A with the individual deletions of C54 or C74 (Figure S4). C28A not only prevented formation of compact folding intermediates, it also increased their heterogeneity. As C28 deletion aggravated folding defects of 54-74 disulfide bond mutants, C28 must have partially compensated for the 54-74 folding defect by participating in oxidative folding. We concluded that C28A in the signal peptide was important for oxidative folding of incompletely folded gp120, most likely for sustaining isomerization of non-native disulfide bonds, and is partially redundant with the 54-74 cysteines for this process.

To analyze whether C28, in addition to redundancy, interacted directly with the 54-74 disulfide bond we used a 110 amino-acid truncation (111X, Figure 3C) for simplicity as it retained the signal peptide and contains a single native cysteine pair. Because formation of this disulfide bond was not detectable by comparing reduced and non-reduced samples, we made use of an alkylation-switch assay [Figure 3D, (Appenzeller-Herzog and Ellgaard, 2008)]. In short, we radiolabeled cells expressing 111X and blocked free cysteines with NEM. Cells then were homogenized and denatured in 2% SDS and incubated again with NEM to block any free cysteines previously shielded by structure. After immunoprecipitation and reduction of disulfide bonds with TCEP, we alkylated resulting free cysteines with mPEG-malemide 5,000, which adds ∼5 kDa of mass for each cysteine alkylated. Samples were immunoprecipitated again to remove mPEG and were analyzed by non-reducing 4-15% SDS-PAGE (Figure 3E).

The 111X construct only showed weak disulfide-bond formation with only ∼13% of molecules forming a disulfide bond (Figure 3E, Wt). Upon removal of the signal-peptide cysteine C28 however, the population that contained a disulfide bond increased significantly to ∼22% (Figure 3F). The presence of C28 thus further destabilized the already unstable 54-74 disulfide bond. The non-native disulfide bond 28-74 in the C54A mutant barely formed, whereas the 28-54 disulfide bond in the C74A mutant was highly variable (Figure 3F). This suggests that disulfide bonds involving the signal-peptide cysteine 28 are unstable and may only occur transiently, a feature consistent with a transient role in disulfide isomerization.

### The N-terminal cysteines form long-range disulfides during early gp160 folding

Gp120 undergoes constant disulfide isomerization during folding (Land et al., 2003) and prolonged association of the signal peptide appears to intrinsically sustain isomerization and may further destabilize the already unstable 54-74 disulfide bond. Moreover, we have shown redundancy of C28 with disulfide bond 54-74, which is why we asked whether the three N-terminal cysteines in gp120 were taking part in long-range disulfide bonds during folding. We removed the V1V2 variable loops, which are not essential for folding and function (Bontjer et al., 2009), and inserted a cleavage site for the protease thrombin through mutagenesis (L125R). This removed all 3 disulfide bonds in V1V2, 126-196 and 131-157 by the loop deletion and 119-205 by mutation (C119-205A). We named the resulting construct gp120Th. Reduction after cleavage produces an N-terminal fragment of ∼15 kDa containing the signal peptide and the 3 N-terminal cysteines C28, C54, and C74, plus a ∼75 kDa fragment containing the rest of gp120 (Figure 4A). If long-range disulfides between the N and C-terminal fragments indeed exist, the cleaved, non-reduced molecule should run in the same position as uncleaved in non-reducing conditions, and should dissociate into the 2 fragments under reducing conditions.

**Figure 4.**
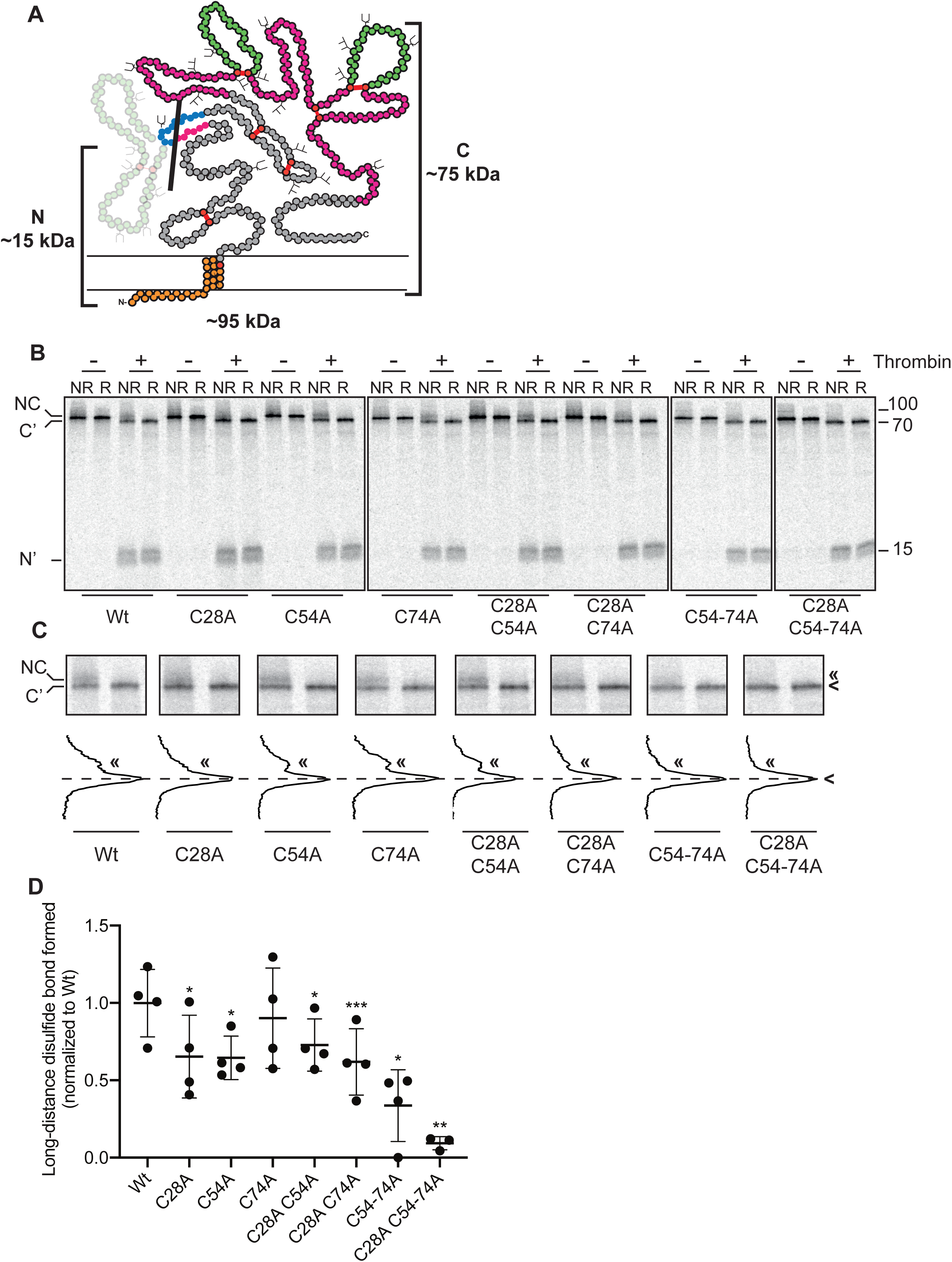
Gp120 exhibits long-distance, non-native disulfide bonds during early folding. **A**) Schematic representation of gp120 thrombin-cleavable construct. Inner domain (grey), outer domain (pink) and variable loops (green) from Figure 1A. Black bar indicates cleavage site for thrombin. **B)** Pulse-chase experiments conducted as in Figure 1B with a 5 min pulse labeling, except that detergent lysates were immunoprecipitated with polyclonal serum HT3. After immunoprecipitation, samples were cleaved with thrombin or mock treated for 15’ at RT. All samples then were analyzed by 15-20% discontinuous SDS-PAGE. NC: non-cleaved, full-length protein, C’: C-terminal fragment; N’: N-terminal fragment. **C)** Zoom in of gels from B showing full-length and C-terminal fragments, lane profiles were generated from autoradiographs in ImageQuant TL. **D)** Quantifications of autoradiographs from **B**. Values were calculated by dividing the signal in the N-terminal fragment by the full-length uncleaved protein and subtracting the value for reducing conditions from non-reducing conditions to determine percent of molecules with a long-distance disulfide bond. Resulting values then were normalized to wild type. Error bars: SD. *: p< 0.05, **: p<0.01, ***: p<0.001. Complete statistical values are listed in Table 1.

Radioactive pulse-chase experiments as described above were modified: instead of deglycosylation with EndoH we denatured the protein with 0.2% SDS and cleaved gp120 with 0.75 U Thrombin. The 2 fragments were separated by 15-20% discontinuous SDS-PAGE. As expected, gp120 that lacked all N-terminal cysteines did not form any long-distance disulfide bonds (Figure 4B and C, C28A C54-74A). We confirmed that wild-type gp120 contained long-distance disulfide bonds between N-terminal cysteines and the rest of the molecule during early folding (Figure 4B and C, Wt). Removal of C28 significantly reduced the number of molecules with a long-range disulfide bond (Figure 4D), likely due to increased stability of the 54-74 disulfide bond in the absence of C28. Strikingly, all cysteine mutants that retained a single cysteine yielded some long-distance disulfides, suggesting that all three N-terminal cysteines could form a non-native pair with downstream cysteines in gp120 (Figure 4B-D).

### Removal of V1/V2 disulfides causes more rapid gp120 folding

Perhaps counterintuitively, the thrombin-cleavage construct (gp120Th) folded faster than full-length gp120 (Figure 5A). Directly after the pulse, gp120Th ran as a more diffuse band whereas full-length gp120 (gp120 Wt) remained close to the reduced-gp120 mobility (Figure 5A NR). This increased compactness shows that gp120Th had already formed more or larger-loop-forming disulfide bonds (Snapp et al., 2017). As a result, signal-peptide cleavage of gp120Th was faster: almost complete for gp120Th after a 1-hour chase, compared to ∼50% cleaved of gp120 Wt (Figure 5A, R).

**Figure 5.**
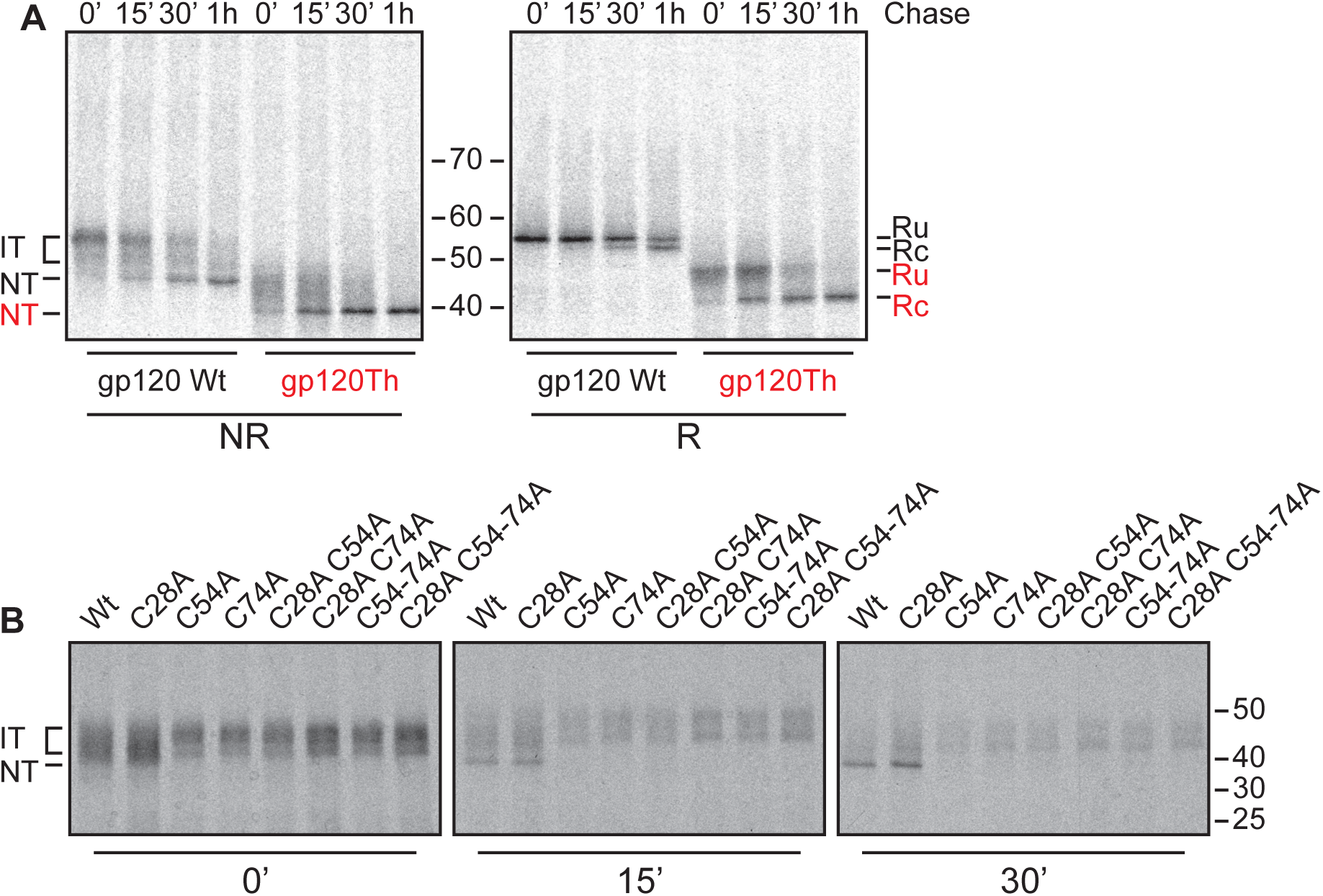
Folding of thrombin-cleavable gp120 construct. **A**) Pulse-chase experiments conducted as in Figure 1B except that HeLa cells were transfected with wild-type full-length gp120 (gp120 Wt) or thrombin-cleavable gp120 (gp120Th) and pulsed for 10 minutes. **B**) Pulse-chase experiments conducted as in Figure 1B except that HeLa cells were transfected with cysteine mutants of gp120Th. Polyclonal serum HT3 was used for immunoprecipitation from detergent lysates and samples were analyzed by 7.5% non-reducing SDS-PAGE. IT: folding intermediates, NT: native gp120, Ru: reduced signal-peptide-uncleaved gp120, Rc: reduced signal-peptide-cleaved gp120. Red text in **(A)** refers to gp120Th running positions.

The 54-74 disulfide-bond mutants in gp120-Wt background fold to a stable intermediate just above the native position [Figures 3B and S4, (van Anken et al., 2008)] whereas the same mutants lacking V1V2 (in gp120Th) failed to accumulate in a single band (Figure 5B), reminiscent of the folding of gp120 C28A C54-74A (Figure 3B). This indicates that V1V2 deletion phenocopied C28 deletion in the 54-74 mutants. Redundancy of V1V2 with C28 was confirmed by the lack of additional effect of C28 removal in the gp120Th 54-74 mutants (Figure 5B).

### C28A results in decreased HIV-1 production and pseudovirus infectivity

As the biochemical data suggested a role for C28 in gp120 folding, we examined the effect of C28A mutation on viral production and infectivity. For this we transfected cells with a molecular clone containing the full HIV genome (LAI strain), containing either wild-type gp160 or the C28A mutant. As the reading frames of *Env* and HIV-1 *Vpu* overlap and mutations in the signal peptide of gp120 can cause changes in the C-terminus of Vpu, we produced the viruses in HEK 293T cells, which are deficient in CD4 and tetherin and therefore do not require Vpu to enhance virus production (Van Damme et al., 2008). We consistently detected significantly less C28A HIV virus than wild-type HIV produced (Figure 6A). Strikingly, the C28A virus was significantly more infectious than wild-type HIV but displayed strong heterogeneity in infectivities, indicative of heterogeneity in C28A gp160 incorporated into the virions (Figure 6B). Due to the severe deficit in virus production, despite increased infectivity, C28A-gp160-containing HIV is not likely to be competitive in nature. Indeed, alignments of >4,300 gp160 sequences from across all subtypes show that C28 is ∼87% conserved (www.hiv.lanl.gov).

**Figure 6.**
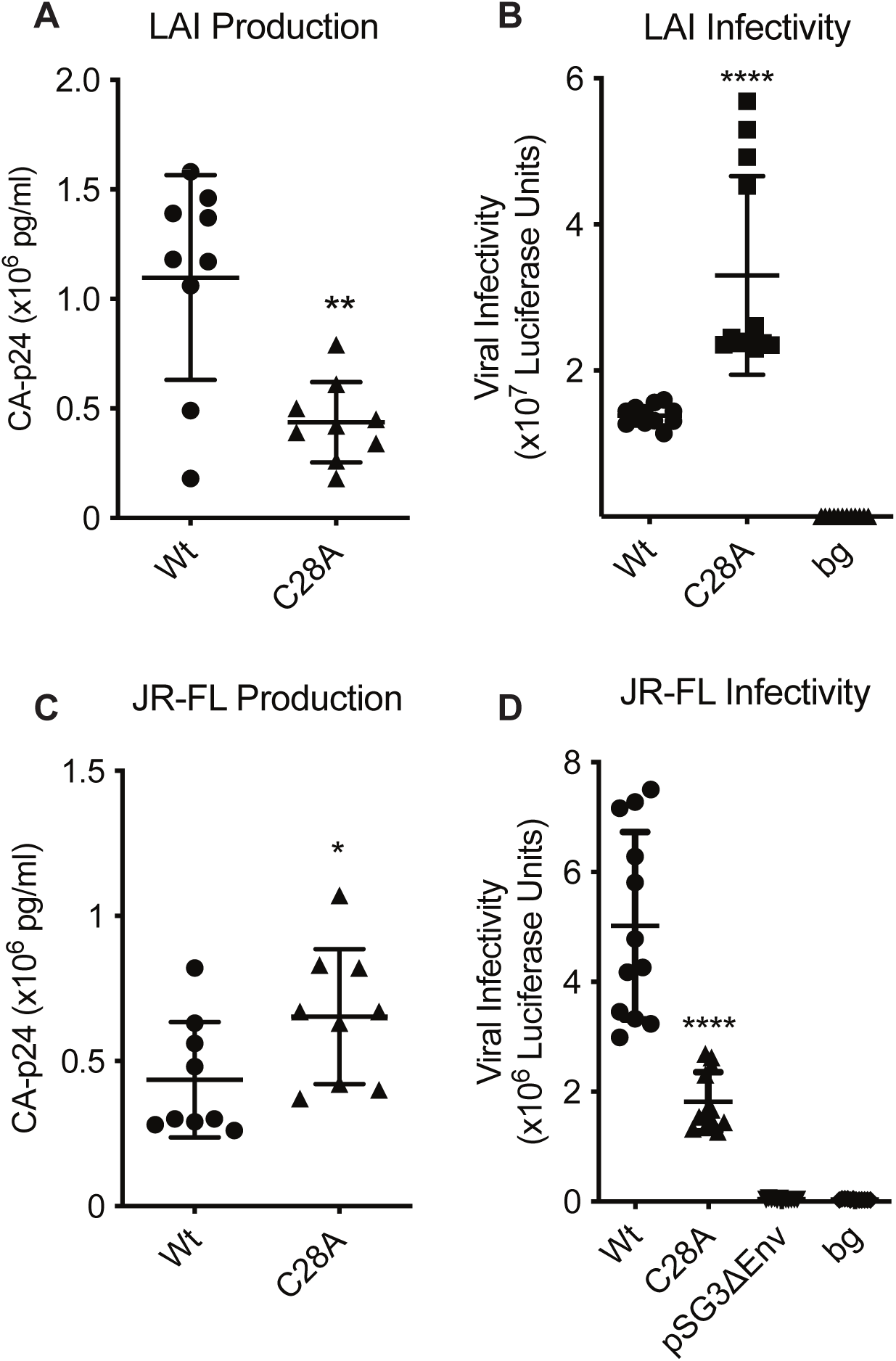
C28A gp160 is detrimental to HIV-1 production and pseudovirus infectivity. **A)** HEK 293T cells were transfected with wild-type or mutant pLAI constructs and virus production was measured by CA-p24 ELISA. **B**) Infection assays were performed as in Figure 2F except with wild-type or C28A gp160 containing HIV-1, as produced in A. bg = background. **C)** Virus produced as in A, except cells expressed wild-type or mutant JR-FL constructs along with packaging plasmids. **D)** Infectivity assays were performed as in Figure 2E. Error bars: SD, pSG3Δ*Env*: virus produced without gp160 plasmid, bg: background, *: p<0.05, **: p<0.01, ****: p<0.0001. Complete statistical values are listed in Table 1.

To uncouple differences in virus production from infectivity we moved to a pseudovirus system, which allows analysis of the effect of C28A gp160 on infectivity alone (Figure 6C and D). As expected, we found very little difference in virus production between wild-type and C28A gp160 (Figure 6C). Infectivity of the C28A gp160 pseudovirus was roughly 60% less than the infectivity of wild type (Figure 6D). We concluded therefore that gp160 conformation, its function, and as a result HIV, suffered from the removal of the signal-peptide cysteine.

## Discussion

During and for 15-30 min after synthesis, the N-terminus of HIV-1 gp160 remains tethered to the ER membrane by its transient signal anchor. We here show that conformational plasticity is enhanced through the cysteine in the signal peptide driving disulfide isomerization, in part via the 54-74 disulfide bond, until the gp120 C-terminus has assembled with the N-terminus, completing the inner-domain β-sandwich and gp120 folding (Figure 7). This triggers signal-peptide cleavage, removing C28 from the protein, halting further isomerization and stabilizing the native gp120 form. This intramolecular quality-control process is essential for viral fitness of HIV and can be impaired and restored by single charge reversals in the gp120 inner domain.

**Figure 7.**
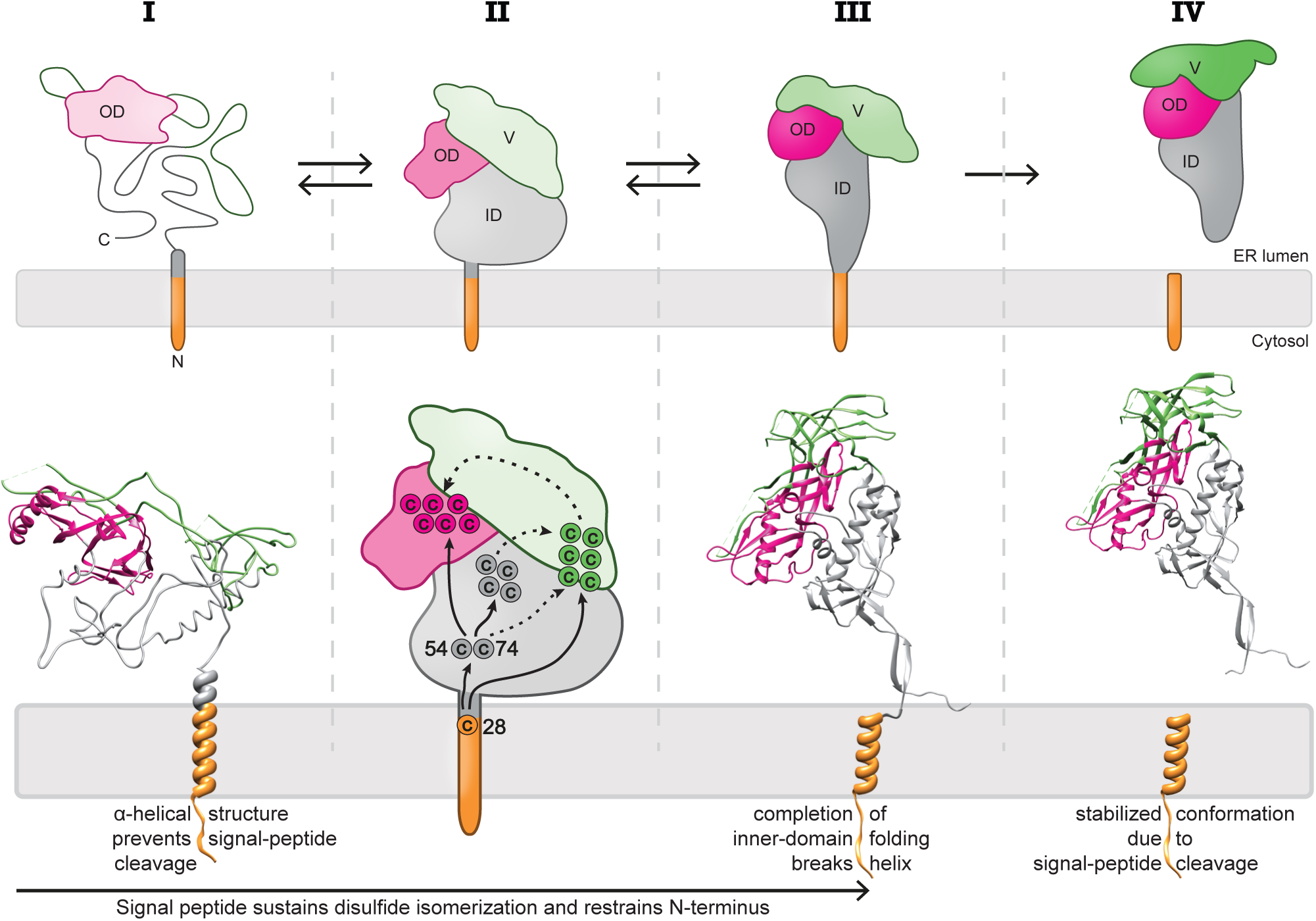
Model for gp120 folding, signal-peptide cleavage and intramolecular disulfide shuffling. **A)** Post-translational domain folding and signal-peptide cleavage of gp120. Grey: inner domain, bright pink: outer domain, green: variable loops, orange: signal peptide, light pink: ribosome, blue: Sec61 translocon. **B)** Conformational changes in the signal peptide and proximal areas during gp120 folding that lead to cleavage. Colors as in **(A)**. **C)** C28 sustains intramolecular disulphide isomerization by interacting with downstream cysteine residues. Solid lines: interactions found experimentally, dashed lines: predicted interactions. Colors as in **(A)**.

### Hierarchy of gp120 folding

The inner domain with the β-sandwich and the outer domain, which contains a stacked double β-barrel, together constitute the minimal folding-capable “core” of gp120 [(Figure 2A), (Garces et al., 2015; Kwong et al., 1998)]. Gp120 is completed with the surrounding variable loops V1V2, V3, and V4 (green in Figures 1A, 2A and B). The core contains six of the nine disulfide bonds in gp120, including the five that are essential for correct folding and signal-peptide cleavage (van Anken et al., 2008). The inner-domain β-sandwich consists of seven strands, six of which are N-terminal. We here show that proper folding of the sandwich requires assembly with the C-terminal β31 strand and formation of the five essential disulfide bonds, which then leads to signal-peptide cleavage [Figure 2A, (van Anken et al., 2008)].

Some folding of the gp120 ’hairpin’ begins during translation and translocation into the ER (Land et al., 2003), but the bulk occurs post-translationally (Figure 7). The (pink) outer domain is the first complete domain to emerge from the translocon and has low contact order, meaning that folding does not require integration of distal residues (Figure 7). It is the first domain to fold, which requires formation of the two native disulfide bonds (296-331 and 385-418) in the β-barrel: gp120 lacking either disulfide bond barely folds past the reduced position in SDS-PAGE (Sanders et al., 2008; van Anken et al., 2008).

The (grey) inner domain of gp120 folds next: individual deletions of its three essential disulfide bonds (54-74, 218-247, and 228-239) fold into more compact structures than the outer-domain deletions (van Anken et al., 2008). The most intriguing is the 54-74 disulfide: the C54-74A mutant accumulates in a sharp band just above the native position, retaining its signal peptide. In contrast, C218-247A and C228-239A fail to form defined intermediates (van Anken et al., 2008), suggesting that these β-sandwich-embracing disulfides (Figure 2A) stabilize the inner domain.

Folding of the inner domain leads to cleavage of the signal peptide. Until that time, the signal peptide acts as signal anchor because it adopts an α-helical conformation that extends past the cleavage site and prevents proteolytic cleavage (Snapp et al., 2017). Folding and integration of the inner domain must break the helix and allow cleavage to occur, as in crystal structures of gp120 this early helical region is a β-strand (Garces et al., 2015). As the disulfide bonds of V1/V2 and gp41 are dispensable for signal-peptide cleavage and their deletion shows no aberration in the folding pathway (van Anken et al., 2008), those domains likely fold after or independent of the outer and inner domains (Figure 7). This is underscored by gp120 folding and signal-peptide cleavage being largely independent of gp41 (Land et al., 2003), and by N- and C-terminal sequences in the gp120 inner domain forming the binding site of gp41 (Garces et al., 2015; Julien et al., 2013; Lyumkis et al., 2013). Gp41 binding may explain apparent inconsistencies between folding and function of some inner-domain mutants (Garces et al., 2015; Yang et al., 2003). The conserved gp41 binding site on the gp120 inner-domain β-barrel also may explain the conservation (and hence value) of the intramolecular quality-control system: it ensures proper folding of this binding site, with high fidelity and well timed, before gp41 folding.

The regulation of signal-peptide cleavage by folding of gp120 implies that the formation of the β-sandwich generates sufficient force to break the attached α-helix and expose the cleavage site. Alpha-helical proteins have lower mechanical strength than β-sheet proteins, which often need to resist dissociation and unfolding; lower mechanical strength facilitates conformational changes to expose transient binding sites or allow signaling (Chen et al., 2015). Only ∼5 pN indeed suffices for exposure of a protease-cleavage site in an α-helix: for proteolytic activation of Notch, cleavage of the NRR domain by ADAM17 (Gordon et al., 2015), and of the talin R3 domain, a 4-helix bundle (del Rio et al., 2009; Yao et al., 2014); the von Willebrand factor A2 domain requires ∼8 pN (Zhang et al., 2009). Pulling apart a β-sheet protein such as Ig domains, OspA, or ubiquitin by shearing needs >160 pN (Brockwell et al., 2003; Carrion-Vazquez et al., 2003; Hertadi et al., 2003), with less force required for peeling (Brockwell et al., 2003). Physiological forces measured so far do not exceed ∼40 pN, a force level at which proteins in general may be destabilized already (Chen et al., 2015). The α-helical region around the cleavage site in gp120 thus would lose the stability competition from the β-sheet in the inner domain, if their structures are incompatible; indeed, in the gp160 structure this α-helical region is a β-strand (Figures 2A and 7). First-time folding, i.e. the completion of the inner-domain β-sandwich by assembly of β31, is likely to generate sufficient force as well, as 7-12 pN allows constant binding of a filamin β-strand to a β-sheet (Rognoni et al., 2012). Formation of the inner-domain disulfide bonds may further raise the stabilizing force (Eckels et al., 2019). The completion of gp120 therefore likely generates the ∼5-pN force needed to break the α-helix and allow the signal peptidase to cleave off the gp120 signal peptide.

### Effects of the attached signal peptide

The postponed cleavage of the signal peptide makes it a transient signal anchor, which acts as membrane tether. This limits conformational freedom and benefits folding, i.e. the integration of the C-terminal β31-strand into the folded inner-domain β-sandwich, as in knotted proteins (Soler and Faisca, 2012).

The prolonged proximity of the free signal-peptide cysteine (28) supports disulfide isomerization and increases conformational plasticity during gp120 folding. Gp120 requires a native set of disulfide bonds to attain its functional 3D structure, but already during synthesis non-native disulfides are formed, which reshuffle over time (Land et al., 2003). The isomerization is detected as “waves” in SDS-PAGE, where the heterogeneous population of folding intermediates oscillates between higher and lower compactness over time [(Figure 3A),(Land et al., 2003)]. Despite extensive isomerization during folding, wild-type gp120 only transiently occupied forms more compact than native. In contrast, the various β-sandwich and uncleavable mutants extensively populated hypercompact states with non-native long-range disulfide bonds, indicating that without stable assembly of the N- and C-termini and resulting retention of the signal peptide, isomerization continues unabated and drives the formation of these hypercompact structures.

The constant disulfide isomerization is sustained by the redox-active cysteine 28 in the signal peptide, as its sulfhydryl group is free to attack existing disulfide bonds. Once gp120 folding has reached a state where isomerization is no longer preferred (N- and C-termini in the inner domain assembled), the signal peptide is cleaved, removing an important, conserved driving force behind isomerization. Cleavage of the signal peptide then acts as a sink because it removes the disulfide-attacking cysteine and pulls the folding equilibrium to the native structure.

### Mode of action of C28

In a short construct, C28 favored a disulfide bond with C54 (Figure 3F). Despite a limited ability to form disulfide bonds with cysteines downstream of the 54-74 bond (Figure 4D), deletion of C28 did not aggravate the C54-74A defective phenotype in the absence of V1V2. C28 hence likely sustains isomerization primarily by constant attack and destabilization of the 54-74 disulfide bond. This propagates a free sulfhydryl group downstream through the folding protein, as the result of an intramolecular electron-transport chain from the more C-terminal cysteines via C54 and C74 to C28 (Figure 7). In the 110-residue gp120 chain (111X), essentially a mimic of a released gp120 nascent chain, in maximally 20% of molecules the 54-74 bond had formed, demonstrating its inherent instability as well as the likelihood that C28 already acts on 54 and 74 during translation.

Only in the presence of V1V2, C28 showed redundancy with the 54-74 disulfide, implying that C28 can fulfill roles otherwise played by C54 and C74 (and vice versa). This suggests that C28 is involved in folding (and isomerization of V1V2) and may play this role by direct interaction with V1V2 cysteines, in absence of C54 and C74, distinct from its 54-74-mediated role in downstream disulfide bonds formation. We cannot exclude that the attack of C28 on the V1V2 disulfides leads to an alternative electron transport chain from C-terminal cysteines via V1V2 to C28. Either way, the redox-active C28 needs to be removed at the end of gp120 folding to ensure stability of the gp120 conformation.

### Intramolecular oxidoreductase and quality control for proper folding

Built-in isomerase activity may seem redundant considering that gp120 folds in the ER, a compartment that contains >20 protein-disulfide-isomerase family members (Jansen et al., 2012). These oxidoreductases are large, bulky proteins however, which cannot catalyze disulfide-bond formation once areas have attained significant tertiary structure. Isomerase activity built into the folding protein allows free-thiol propagation (in essence electron transport) in areas otherwise unreachable by folding enzymes. This perhaps should not be too surprising given that in folded proteins the majority of cysteines are solvent inaccessible (Srinivasan et al., 1990) and during folding disulfide bonds become resistant to reduction with DTT (Tatu et al., 1993; Tatu et al., 1995).

An example of intramolecular disulfide isomerization is the cysteine in the pro-peptide of bovine pancreatic trypsin inhibitor (BPTI), which increased both the rate and yield of BPTI folding (Weissman and Kim, 1992). The majority of disulfide formation during *in-vitro* folding of BPTI results from intramolecular disulfide rearrangements (Creighton et al., 1993; Darby et al., 1995; Weissman and Kim, 1995). Transfer of free thiols between lumenal and transmembrane domains in the ER has been demonstrated for vitamin-K-epoxide reductase (Liu et al., 2014; Schulman et al., 2010), indicating that such exchanges are possible. While C28 is located in the transmembrane α-helix (Snapp et al., 2017), suggesting immersion in the membrane, sliding of transmembrane domains up and down in the membrane is possible (Borochov and Shinitzky, 1976; Danielson et al., 1994; Mowbray and Koshland, 1987). As C28 is part of the consensus sequence for signal-peptide cleavage, it likely is exposed to the ER lumen at least part of the time.

Not only intramolecular oxidoreductase activity offers an advantage, also intramolecular quality control. Release of a protein from the ER requires its folding to the extent that chaperones do not bind anymore, for instance due to shielding of hydrophobic residues from Hsp70 chaperones. The intramolecular quality control we here describe ensures a much more subtle regulation of conformational quality. Gp160’s function as HIV-1 fusion protein requires the native interaction between gp41 and the gp120 inner domain. Only proper exposure of the gp41 binding site in gp120 will lead to a functional protein. Intramolecular quality control ensures precision to the level of single residues as well as precision of timing.

### Conserved and multiple roles for signal peptides

Post-translational signal-peptide cleavage of gp120 is conserved across different subtypes of HIV-1 as biochemical properties, even if sequences are not strictly conserved (Snapp et al., 2017). This appears to be more general, as in other organisms signal peptides mutate at a lower rate than the surrounding mature peptide (Morrison et al., 2003; Williams et al., 2000), or they mutate at the same rate, but with an increased proportion of null (Veitia and Caburet, 2009) or function-preserving mutants (Garcia-Maroto et al., 1991). Function-altering mutants often have deleterious effects (Bonfanti et al., 2009; Piersma et al., 2006).

Detailed kinetic analysis of signal-peptide cleavage has not been reported for a great number of proteins, and Western blotting often does not offer the necessary resolution, but gp160 is not alone in its biosynthesis-dependent and biosynthesis-regulated signal-peptide cleavage (Anjos et al., 2002; Daniels et al., 2003; Matczuk et al., 2013; Rutkowski et al., 2003; Zschenker et al., 2001). Whereas structure regulates cleavage in HCMV US11 (Rehm et al., 2001; Tamura et al., 2011), function regulates cleavage in ERAD-associated protein EDEM1 (Rehm et al., 2001; Tamura et al., 2011). These studies demonstrate that a variety of conditions including nascent-chain length and N-glycan addition can play a role in signal peptide cleavage. Signal peptides are more than address labels and folding and signal-peptide cleavage are more interdependent than originally thought (Li et al., 1994; Rehm et al., 2001; Tamura et al., 2011).

We argue that late signal-peptide cleavage may be much more common than biochemical experiments have uncovered. Cleavage may occur at any time from co-translationally until late post-translationally. Considering the low rate of protein synthesis, ∼3 to 6 amino acids per second (Braakman et al., 1991; Horwitz et al., 1969; Ingolia et al., 2011; Knopf and Lamfrom, 1965), it can lead to long average synthesis times (∼1.5–3 min for gp120 and ∼2–5 min for gp160). In fact, translation rates are much more heterogeneous (Ingolia et al., 2019): nascent chains of influenza virus HA may take >15 min to complete, corresponding to a rate of less than one residue per second [(Braakman et al., 1991); unpublished observations]. For large proteins, the difference between early and late co-translational cleavage leaves a window of several minutes, during which the signal peptide functions as an anchor tethering the protein to the ER membrane. Sequence features in the signal peptide, such as an exposed cysteine, are given the opportunity to interact with the folding protein.

The membrane tether limits conformational freedom of the protein and reduces overall conformational entropy, which is predicted to increase fidelity of protein folding and stability (Dill and Alonso, 1988; Zhou, 2008; Zhou and Dill, 2001). This may well benefit the formation of N- and C-terminal contacts in proteins, which are present in ∼50% of soluble PDB structures (Krishna and Englander, 2005) and are present in multiple multimeric viral glycoproteins (Chen et al., 1998; Garces et al., 2015; Gogala et al., 2014; Sauter et al., 1992; Sun et al., 2014).

Here we have presented compelling evidence for the direct functional contribution of the signal peptide to HIV-1 gp160 folding. The signal peptide drives disulfide isomerization of gp120 during folding, increasing conformational plasticity while tethering the N-terminus, and functions as quality control organizer, leaving only after near-native conformation has been attained. As evidence grows, it becomes clear that signal peptides demonstrate functions far beyond their originally assigned roles as cellular postal codes.

## Acknowledgements

We would like to thank members of the Braakman-Van der Sluijs and Sanders labs for their fruitful discussions and insights. In particular Peter van der Sluijs for critical reading of the manuscript and Joseline Houwman for critical reading of the manuscript and design of figure 7. This work was supported by grants from the Dutch Research Council (NWO)- Chemical Sciences (I.Br, N.M, A.L, M.Q), the European Union 7th framework program, ITN “Virus Entry” (I.Br, N.M, M.Q), the European Union’s Horizon 2020 research and innovation program under grant agreement No. 681137 (R.W.S. and I.Bo). R.W.S. is a recipient of a Vici grant from the Dutch Research Council (NWO).

## Author Contributions

Conceptualization: N.M, M.Q and I.B. Methodology: N.M. Investigation: N.M, M.Q, I.Bo and A.L. Writing – Original Draft: N.M, M.Q, and I.B. Writing – Review & Editing: N.M, M.Q, I.Bo, R.W.S, A.L and I.B. Funding Acquisition: R.W.S and I.B.

## Declaration of Interests

The authors declare no competing interests.

## Materials and methods

### Plasmids, antibodies, reagents and viruses

The full-length molecular clone of HIV-1_LAI_ (pLAI) was the source of wild-type and mutant viruses (Peden et al., 1991). The QuikChange Site-Directed Mutagenesis kit (Stratagene) was used to introduce mutations into *Env* in plasmid pRS1 as described before; the entire *Env* gene was verified by DNA sequencing (Sanders et al., 2004). Mutant *Env* genes from pRS1 were cloned back into pLAI as SalI-BamHI fragments. For transient transfection of gp120/160 we used the previously described pMQ plasmid (Snapp et al., 2017). C-terminal truncations were generated by PCR of wt gp120 and Gibson assembled back into XbaI/XhoI digested pMQ. The thrombin-cleavable construct was designed based on stable V1V2 loop deletion number 2 (Bontjer et al., 2009) and generated from gp120 C119-205A using Gibson assembly (Gibson et al., 2009). All point mutations were introduced using QuikChange Site-Directed mutagenesis as above.

For immunoprecipitation: we used the previously described polyclonal rabbit anti-gp160 antibody 40336 which recognizes all forms of gp120 (Land et al., 2003), polyclonal antibody HT3 (NIH531) which was obtained from the NIH AIDS reagent program and α-HA tag antibody “MrBrown” produced by us (Schildknegt et al., 2019).

Although we studied gp160 of the LAI isolate, we followed the canonical HXB2 residue numbering (GenBank: K03455.1), which relates to the LAI numbering as follows: because of an insertion of five residues in the V1 loop of LAI gp160, all cysteine residues beyond this loop have a number that is 5 residues lower in HXB2 than in LAI: until Cys131, numbering is identical, but Cys162 in LAI becomes 157 in HXB2, etc.

Thrombin was purchased as a lyophilized power from Sigma Aldrich (T-6634) and stored in thrombin-storage buffer [50 mM Sodium Citrate pH 6.5, 200 mM NaCl, 0.1% BSA (w/v), 50% glycerol (w/v)].

### Cells and transfections

The SupT1 cell line was cultured in Advanced RPMI 1640 medium (Gibco), supplemented with 1% fetal calf serum (v/v, FCS), 2 mM L-glutamine (Gibco), 15 units/ml penicillin and 15 µg/ml streptomycin. The TZM-bl reporter cell line, obtained from NIH AIDS Research and Reference Reagent Program, Division of AIDS, NIAID, NIH (John C. Kappes, Xiaoyun Wu, and Tranzyme, Inc., (Durham, NC)), the HEK293T cell line, and the C33A cell line were cultured in Dulbecco’s modified Eagle medium (Gibco) containing 10% FCS, 100 units/ml penicillin and 100 µg/ml streptomycin. HeLa cells (ATCC) were maintained in MEM containing 10% FCS, nonessential amino acids, glutamax and penicillin/streptomycin (100 U/ml). Twenty-four hours before pulse labeling, HeLa cells were transfected with pMQ gp120/gp160 or HA constructs using polyethylenimine (Polysciences) as described before (Hoelen et al., 2010).

### Virus production

Virus stocks were produced by transfecting HEK293T cells with wild-type or mutant pLAI constructs using the Lipofectamine 2000 Transfection Reagent (Invitrogen) per manufacturer’s protocol. Production of virus stocks on C33A cells were done by calcium-phosphate precipitation. The virus-containing culture supernatants were harvested 2 days post-transfection, stored at -80°C, and the virus concentrations were quantitated by CA-p24 ELISA as described before (Moore and Jarrett, 1988). These values were used to normalize the amount of virus used in subsequent infection experiments.

### Single Cycle Infection

The TZM-bl reporter cell line stably expresses high levels of CD4 and HIV-1 coreceptors CCR5 and CXCR4 and contains the luciferase and β-galactosidase genes under the control of the HIV-1 long-terminal-repeat (LTR) promoter (Wei et al., 2002). Single-cycle infectivity assays were performed as described before (Bontjer et al., 2009; Bontjer et al., 2010). In brief, one day prior to infection, 17 x 10^6^ TZM-bl cells per well were plated on a 96-well plate in DMEM containing 10% FCS, 100 units/ml penicillin and 100 µg/ml streptomycin and incubated at 37°C with 5% CO_2_. A fixed amount of virus LAI virus (500 pg of CA-p24) or a fixed amount of JR-FL or LAI pseudo-virus (1,000 pg of CA-p24) was added to the cells (70-80% confluency) in the presence of 400 nM saquinavir (Roche) to block secondary rounds of infection and 40 µg/ml DEAE in a total volume of 200 µl. Two days post-infection, medium was removed, cells were washed with phosphate-buffered saline (50 mM sodium phosphate buffer, pH 7.0, 150 mM NaCl) and lysed in Reporter Lysis buffer (Promega). Luciferase activity was measured using a Luciferase Assay kit (Promega) and a Glomax luminometer (Turner BioSystems) per manufacturer’s instructions. Uninfected cells were used to correct for background luciferase activity. All infections were performed in quadruplicate.

### Folding assay

HeLa cells transfected with wild-type or mutant gp160/gp120 constructs were subjected to pulse-chase analysis as described before (McCaul et al., 2019; Snapp et al., 2017). In short, cells were starved for cysteine and methionine for 15-30 min and pulse labeled for 5 min with 55 µCi/ 35-mm dish of Easytag express ^35^S protein labeling mix (Perkin Elmer). Where indicated (+DTT), cells were incubated with 5 mM DTT for 5 min before and during the pulse. The pulse was stopped, and chase started by the first of 2 washes with chase medium containing an excess of unlabeled cysteine and methionine. At the end of each chase, medium was collected, and cells were cooled on ice and further disulfide bond formation and isomerization was blocked with 20 mM iodoacetamide. Cells were lysed and detergent lysates and medium samples were subjected to overnight immunoprecipitation at 4°C with polyclonal antibody 40336 against gp160.

### Deglycosylation, SDS-PAGE, and autoradiography

Where appropriate, to identify gp160 folding intermediates, glycans were removed from lysate-derived gp120 or gp160 with Endoglycosidase H (Roche) treatment of the immunoprecipitates as described before (Land et al., 2003).

Samples were subjected to non-reducing and reducing (25 mM DTT) SDS-PAGE. Gels were dried and exposed to super-resolution phosphor screens (FujiFilm) or Kodak Biomax MR films (Carestream). Phosphor screens were scanned with a Typhoon FLA-7000 scanner (GE Healthcare Life Sciences). Quantifications were performed with ImageQuantTL software (GE Healthcare Life Sciences).

### mPEG treatment

HEK 293T cells transfected with wild-type or mutant 111X were subjected to radioactive labeling as described above. At the end of the labeling, cells were transferred to ice and incubated in Dulbecco’s PBS without Ca^2+^ and Mg^2+^ containing 20 mM *N*-ethyl malemide (NEM) and 5 mM EDTA. Cells then were subjected to a modified “double-alkylation variant” mPEG treatment as described by Appenzeller-Herzog and Ellgaard (Appenzeller-Herzog and Ellgaard, 2008). In short, cells were homogenized by passage through a 25-G needle and proteins denatured with 2% SDS for 1 h @ 95 °C. Samples then were alkylated again with 20 mM NEM before immunoprecipitation with anti-HA tag antibody MrBrown for 2 hours at 4 °C. After immunoprecipitation, samples were denatured and reduced with 25 mM TCEP followed by incubation with 15 mM mPEG-mal 5000 for 1 h at room temperature. Samples were immunoprecipitated again via the HA-tag and analyzed by 4-15% non-reducing gradient SDS-PAGE (BioRad) and processed as before.

### Thrombin cleavage

After HeLa cells transfected with various thrombin-cleavable constructs were pulse-labeled as described above, detergent lysates were immunoprecipitated with antibody HT3 for 1 h at 4°C with rotation. Immunoprecipitates were washed and resuspended in 15 µl thrombin cleavage buffer (20 mM Tris-HCl, pH 8.4, 150 mM NaCl, 2.5 mM CaCl_2_) + 0.2% SDS and denatured for 5 minutes at 95°C. SDS was quenched by addition of 10 µl cleavage buffer + 2% Tx100. Thrombin (0.75 U) in 5 µl cleavage buffer then was added to samples and incubated for exactly 15 minutes. For mock-digested samples, an equivalent volume of thrombin storage buffer was added instead. Digestion was stopped by the addition of hot (95°C) 5X sample buffer and immediately placing in a 95°C heat block for 5 minutes. Samples then were subjected to non-reducing or reducing (25 mM DTT) 15-20% discontinuous-gradient SDS-PAGE and processed as before.

### Statistical Reporting

Statistics for each experiment were calculated using Prism 7 (Graphpad). For experiments in Figures 2D-F differences were assessed using a one-way ANOVA with follow-up testing to analyze differences between specific pairs with p values corrected for multiple comparisons. For experiments in Figure 3F and 4D differences were assessed using paired t-tests between wild-type and mutants. For experiments in Figure 6A-D differences were assessed using unpaired t-tests. A complete list of all pairs examined, statistical methods and resulting p values can be found in Table 1.

**Figure S1. N-terminal truncations of gp120 retain their signal peptides.**

Pulse-chase experiments were performed as in Figure 1B except that HeLa cells were transfected with the indicated gp120 truncations. Detergent lysates were immunoprecipitated either with polyclonal serum 40336 (a) or a polyclonal serum that recognizes the signal peptide (b).

**Figure S2. Inner domain β-sandwich mutants affect gp120 folding and HIV infectivity.**

**A)** Pulse-chase experiments were performed as in Figure 2C except that HeLa cells were transfected with the indicated mutants. **B)** Uncropped gels from Figure 2C. IT: folding intermediates, NT: native gp120, Ru: reduced signal-peptide-uncleaved gp120, Rc: reduced signal-peptide-cleaved gp120. **C)** Quantifications performed as in Figure 2D. **D)** Quantifications performed as in Figure 2E. **E)** Infection assays were performed as in Figure 2F. Error bars: SD.

**Figure S3. Synchronized folding of gp120 Wt and C28A.**

**A)** Pulse-chase experiment was performed as in Figure 1B except that cells expressing Wt or C28A gp120 were treated from 5 minutes before the pulse with 2 mM puromycin and chased in the presence of 500 mM cycloheximide. Samples were analyzed by reducing 7.5% SDS-PAGE after immunoprecipitation. **B)** Lane profiles from **A.** Ru: reduced signal-peptide-uncleaved gp120, Rc: reduced signal-peptide-cleaved gp120.

**Figure S4. Removal of signal-peptide cysteine C28 aggravates folding phenotype of 54-74 disulfide-bond mutants**

Pulse-chase experiments were performed as in Figure 1B except that HeLa cells expressed C28 and disulfide-bond 54-74 mutants.

## Notes

### Competing Interest Statement

The authors have declared no competing interest.

